# Imperceptible gamma-band sensory stimulation enhances episodic memory retrieval

**DOI:** 10.1101/2023.07.21.550057

**Authors:** Benjamin J. Griffiths, Daniel E. Weinert, Ole Jensen, Tobias Staudigl

**Author notes:** Corresponding author: Benjamin J. Griffiths. **Funding statement:** B.J.G. is funded by a Leverhulme Trust Early Career Fellowship (ECF-2021-628). **Data/code availability:** All data and code relating to this paper can be found on Github (https://github.com/benjaminGriffiths/harmonic-flicker). The project pre-registration can be found on the Open Science Framework (https://osf.io/qkm2g/). **Supplementary materials:** For the sake of convenience, all supplementary materials have been appended to this document and can be found after the reference section.

## Abstract

Enhanced gamma activity (30-100Hz) coincides with the successful recall of episodic memories, but it remains unknown whether this oscillatory activity is a cause or a consequence of the retrieval process. To address this question, we asked human participants to complete a paired associates memory task while undergoing sensory stimulation (at 65Hz, 43.3Hz and 32.5Hz). We observed that 65Hz and 32.5Hz sensory stimulation enhances recall compared to a baseline condition without stimulation. No similar effect was observed following 43.3Hz stimulation. Notably, while almost all participants could perceive 32.5Hz and 43.3Hz sensory stimulation, only a small proportion of participants (∼10%) could perceive the 65Hz visual flicker, suggesting 65Hz sensory stimulation acts as an imperceptible intervention to enhance recall. To understand the dual action of 65Hz and 32.5Hz sensory stimulation on recall, we built three pyramidal-interneuronal network gamma (PING) models and drove them using the same stimulation protocols as in the behavioural task. The behavioural results could be reproduced by stimulating an endogenous ∼32Hz oscillation, but not by stimulating an endogenous ∼65Hz oscillation nor by stimulating a network without an endogenous oscillation. These results suggest that imperceptible 65Hz sensory stimulation enhances recall by harmonically entraining an endogenous ∼32.5Hz oscillation. Based on these findings, we propose that “slow” gamma oscillations play a causal role in episodic memory retrieval.

## Introduction

In humans, extensive research demonstrates that episodic memory formation and retrieval are both accompanied by an increase in gamma-band activity (30-100Hz) ^e.g. 1–45^. Interestingly, growing evidence suggests that the canonical gamma band can be divided into a least two parts, with “fast” gamma oscillations (60-100Hz) supporting encoding and “slow” gamma oscillations (20-40Hz) supporting retrieval ^7, 29, 46–48^. However, despite numerous demonstrations of these brain-behaviour correlations, it is unclear whether gamma oscillations play a causal role in episodic memory. Here, we set out to address this question using visual sensory stimulation.

Visual sensory stimulation is a technique in which a stimulus’s luminance oscillates over time and aims to modulate ongoing neural activity. While low-frequency sensory stimulation can modulate endogenous low-frequency oscillations^49^ and enhance episodic memory formation^50^, and gamma-band sensory stimulation can induce gamma-band activity in the thalamus, visual cortex and hippocampus^51–55^, it remains unclear whether gamma-band sensory stimulation can enhance recall performance above and beyond what is observed when stimulation is not applied. Importantly, as sensory stimulation using fast gamma rhythms (>60Hz) involves flickering the stimuli at rates above the flicker fusion threshold, the flickers are imperceptible to the naked eye and, therefore, may provide an imperceptible intervention to enhance episodic memory.

This is a paper of two parts. First, we ask whether gamma-band sensory stimulation can enhance memory performance in human participants. By analysing memory performance in a paired-associates task with a series of logistic regression models, we find support for this idea. Specifically, we demonstrate that memory performance can be enhanced by delivering either 32.5Hz or 65Hz sensory stimulation during recall. Second, we explore why 32.5Hz and 65Hz sensory stimulation exerts dual effects on recall, with a particular interest in addressing the apparent contradiction of “fast” gamma stimulation aiding retrieval (rather than encoding ^e.g., 29, 46, 48, 56^). In a collection of computational models, we demonstrate that, for both 32.5Hz and 65Hz sensory stimulation, the enhancement of recall is best explained by the entrainment of an endogenous, “slow” gamma (∼32.5Hz) oscillation. These results hint that 65Hz sensory stimulation can act as an imperceptible intervention to aid episodic memory retrieval by modulating an endogenous “slow” gamma oscillation.

## Results

### 65Hz and 32.5Hz sensory stimulation delivered during retrieval enhances memory recall

In the first behavioural experiment, participants completed a paired associates task where visual sensory stimulation was delivered during both memory formation and retrieval (see figure 1A). We used stimulation frequencies of 65Hz, 43.3Hz, and 32.5Hz, and included an additional baseline condition that involved no visual flicker. Importantly, the stimulation frequencies presented at encoding and retrieval were counterbalanced such that each combination of frequencies [e.g., 65Hz at encoding and 43.3Hz at retrieval; 65Hz at encoding and no stimulation at retrieval] was encountered with equal regularity, allowing for the effects of stimulation on retrieval to be distinguished from the effects of stimulation on encoding. Offline, we used logistic regression models to assess the impact of sensory stimulation on memory performance while controlling for potential confounds including response times, experiment duration, and the stimulation frequency used at the other memory stage (see figure 1B). After pooling beta weights across participants, we then used one-sample t-tests (controlling for multiple comparisons using Bonferroni correction) to assess whether sensory stimulation led to notable changes in memory performance from baseline.

**Figure 1.**
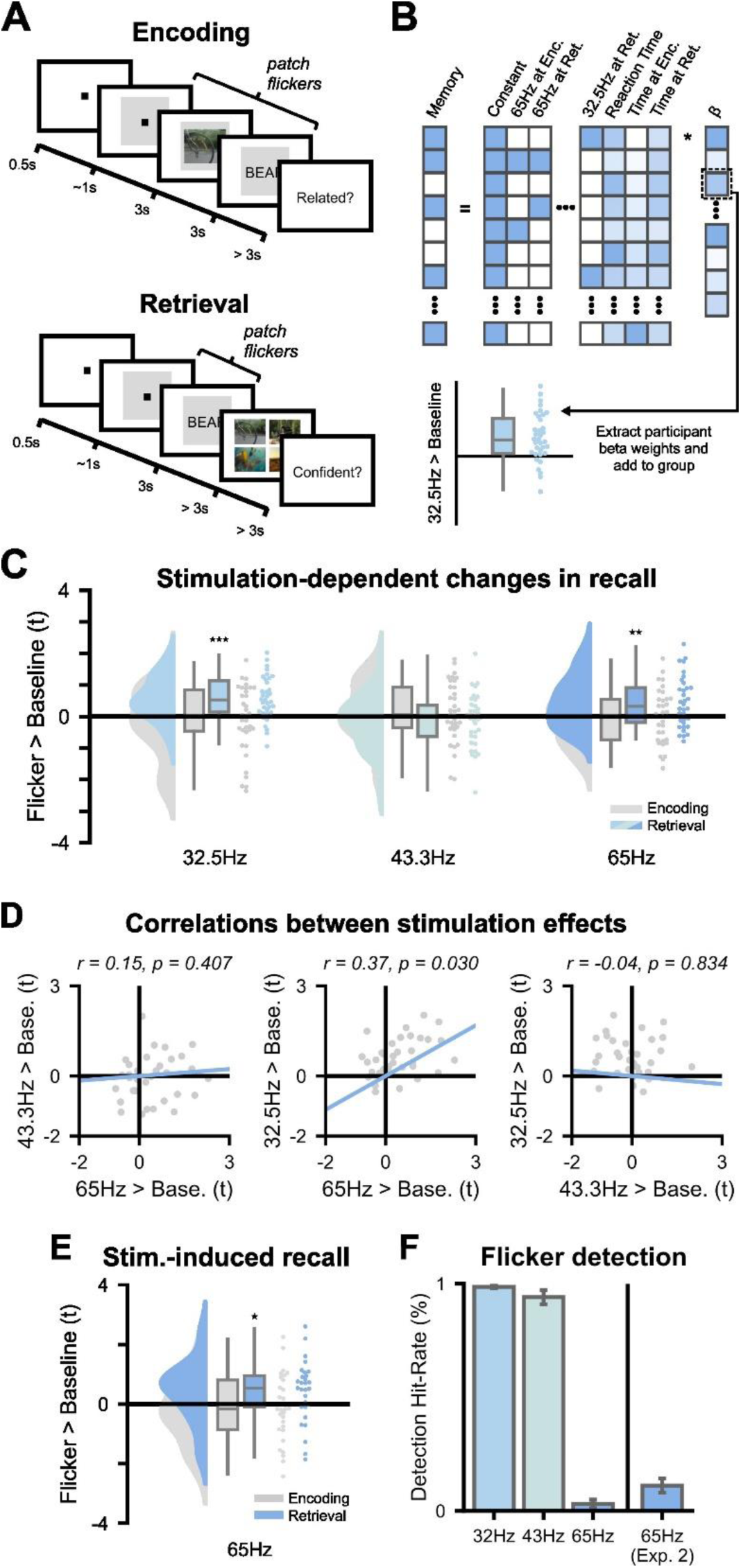
65Hz and 32.5Hz sensory stimulation modulates memory performance. **(A)** Main experimental paradigm. Participants learned video-word pairs and, after a brief distractor task, were asked to recall the video using the word as a cue. Critically, during both encoding and retrieval, the screen either flickered at 32.5Hz, 43.3Hz or 65Hz, or did not flicker. The flickering protocol was counterbalanced such that the flicker at encoding was in no way predictive of the flicker that would occur at retrieval. Logistic regression models were used to delineate the effect of encoding stimulation from the effect of retrieval stimulation. **(B)** Depiction of statistical approach. Following the pre-registered approach, we ran participant-specific logistic regressions to estimate the impact of different stimulation frequencies on memory performance whilst controlling for confounds such as response time and fatigue (modelled as the time elapsed). **(C)** Raincloud plots depicting how 65Hz and 32.5Hz sensory stimulation enhanced recall specifically during retrieval. The y-axis measures t-values returned from the participant-specific logistic models. The x-axis refers to the different stimulation conditions. Grey plots depict the impact of encoding stimulation on recall performance. Coloured plots depict the impact of retrieval stimulation on recall performance. Boxplots depict the median, interquartile range, and minima/maxima of the data. Individual dots depict individual participants. **(D)** Correlation between retrieval stimulation effects for different frequencies. Individual dots depict individual participants. The blue line reflects the line-of-best-fit. Participants which benefited from 32.5Hz stimulation also benefited from 65Hz stimulation. **(E)** Raincloud plots depicting a replication of how 65Hz sensory stimulation enhanced recall specifically during retrieval. All plot details match those described in figure 1B. **(F)** The detectability of different frequencies of sensory stimulation. The y-axis measures the percentage of times participants detected the flicker. The x-axis refers to the different flicker frequencies. While the slower flickers (32.5/43.3Hz) were easily detected, the 65Hz flicker could only be detected by a small minority of participants.

One-sample t-tests revealed that 65Hz visual stimulation during retrieval enhanced memory performance relative to when no stimulation was applied (t(33) = 3.22, Cohen’s d*_z_* = 0.56, p_bonf_ = 0.017; see figure 1C). A similar effect was observed following 32.5Hz sensory stimulation (t(33) = 5.08, Cohen’s d_z_ = 0.87, p_bonf_ < 0.001). Sensory stimulation at 43.3Hz led to no significant change in memory performance relative to baseline (t(34) = -0.84, Cohen’s d_z_ = -0.14, p_bonf_ > 0.5). A “two one-sided test” (TOST) procedure assuming a medium effect size (Cohen’s d_z_ = 0.5; marginally smaller than the 32.5/65Hz effect sizes above) suggested that memory performance following 43.3Hz visual stimulation at retrieval was equivalent to when no sensory stimulation occurred (t_upper_(34) = 2.08, t_lower_(34) = -3.75, p = 0.023). Sensory stimulation delivered during encoding had no influence on subsequent memory (65Hz: t(34) = -0.07, Cohen’s d_z_ = -0.01, p_bonf_ > 0.5; 43.3Hz: t(34) = 0.85, Cohen’s d_z_ = 0.15, p_bonf_ > 0.5; 32.5Hz: t(34) = -0.55, Cohen’s d_z_ = -0.10, p_bonf_ > 0.5). The TOST procedure arrived at the same conclusion: memory performance for all stimulation conditions were equivalent to when no stimulation was applied (65Hz: t_upper_(33) = 2.85, t_lower_(33) = -2.98, p = 0.004; 43.3Hz: t_upper_(34) = 3.77, t_lower_(34) = -2.06, p = 0.024; 32.5Hz: t_upper_(34) = 2.36, t_lower_(34) = -3.74, p = 0.012). These results suggest that 32.5Hz and 65Hz visual stimulation delivered during retrieval boosts recall.

Simplifying the analysis pipeline and focusing solely on the percentage change in memory performance produced the same results. Sensory stimulation at 32.5Hz and 65Hz led to, on average, a ∼10.1% and ∼8.0% increase (respectively) in pairs recalled relative to the baseline condition. These differences were statistically significant (32.5Hz: t(33) = 3.95, Cohen’s d_z_ = 0.69, p = 0.001; 65Hz: t(32) = 3.65, Cohen’s d_z_ = 0.64, p = 0.002). Sensory stimulation at 43.3Hz, on the other hand, led to a ∼1.3% decrease in pairs recalled relative to baseline. This difference was not significant (t(33) = -0.63, Cohen’s d_z_ = -0.11, p > 0.5; TOST: t_upper_(33) = 2.24, t_lower_(33) = -3.50, p = 0.015). These results complement our initial results by demonstrating that 32.5Hz and 65Hz sensory stimulation applied during memory retrieval enhances memory performance by 8-10%.

Notably, the beneficial effects of 32.5Hz and 65Hz sensory stimulation overlapped: Participants who benefited from the 32.5Hz stimulation also benefited from the 65Hz stimulation (Pearson’s r = 0.37, p = 0.030; see figure 1D). Neither 32.5Hz nor 65Hz stimulation effects correlated with 43.3Hz stimulation effects (32.5-43.3Hz correlation: Pearson’s r = -0.04, p = 0.834; 65-43.3Hz correlation: Pearson’s r = 0.15, p = 0.407). These results suggest that the beneficial effects of 32.5Hz and 65Hz sensory stimulation are linked to a common phenomenon.

To test the visual detection of the flicker, participants completed a perception task in which they were shown a video-word sequence (identical to the encoding stage of the task) while the display either flickered at one of the stimulation frequencies or did not flicker. After each video-word sequence, participants had to report whether they thought the screen was flickering. Of the 34 participants who completed this task, all participants could reliably detect the 32.5Hz luminance change, 33 of the 34 participants could detect the 43.3Hz luminance change, but only one participant could detect the 65Hz luminance change to a degree greater than what would be expected by chance (see figure 1E). These results suggest that 65Hz visual sensory stimulation is imperceivable to most participants.

Importantly, we could replicate the memory-boosting effect of 65Hz sensory stimulation in a second sample. The experiment matched the first task, except that only 65Hz sensory stimulation and the no stimulation baseline condition were included. Aligning with the results of the first experiment, 65Hz visual stimulation deployed at the moment of recall enhanced retrieval success relative to when no stimulation was applied (t(28) = 2.21, Cohen’s d_z_ = 0.42, p = 0.018; see figure 1F). The protocol led to a ∼5.2% increase in performance relative to baseline (a difference across participants that was significantly greater than chance: t(28) = 2.39, Cohen’s d_z_ = 0.45, p = 0.012). No similar effect was observed for encoding-based 65Hz stimulation (t(28) = -0.62, Cohen’s d_z_ = -0.12, p > 0.5), with a TOST procedure suggesting memory performance was equivalent between the 65Hz and baseline conditions (t_upper_(28) = 2.02, t_lower_(28) = -3.27, p = 0.026). In the flicker perception task, 6 of 36 participants (∼11.1%) were able to perceive the flicker, again suggesting that most participants could not perceive 65Hz sensory stimulation (resulting in ∼10% of participants being able to perceive the 65 Hz flicker across the two experiments).

The observation that stimulation did not benefit encoding suggests that stimulation does not aid retrieval by enhancing perception/attention because, under a hypothesis which stipulates that stimulation does indeed enhance perception/attention, such an effect should also facilitate encoding. An additional control experiment provides further support to suggest that stimulation aids retrieval-specific processes (as opposed to more general perceptual/attentional/motor effects; see supplementary figure 1).

Taken together, the results presented here suggest that visual 32.5Hz and 65Hz sensory stimulation can aid the recall of episodic memories, quite possibly through aiding the reactivation/reinstatement of the episodic memory trace. Of course, this begs the question “how?”. In the next section, we aim to answer this by exploring the effects of harmonic entrainment on a series of computational models.

### Harmonic entrainment of an endogenous “slow” gamma oscillation explains the beneficial effects of 32.5Hz and 65Hz sensory stimulation on memory retrieval

In this section, we ask whether the observed behavioural effects can be explained by oscillatory entrainment (that is, the strengthening of an endogenous rhythm) and, if so, what frequency this endogenous oscillation takes. In particular, we focus on the concept of “harmonic entrainment”, which refers to the phenomenon in which an exogenous input oscillating at a harmonic of the endogenous oscillatory frequency can entrain the endogenous frequency. As several studies have already demonstrated that harmonic entrainment can arise *in silico* and *in vivo* ^57–60^, our model is not intended to give a comprehensive account of the phenomenon, but instead aims to determine whether such entrainment could explain the stimulation-related changes in memory performance reported above and, more generally, whether harmonic entrainment can bring about behavioural change.

To this end, we built three neural networks of Izhikevich^61^ regular spiking neurons and fast-spiking interneurons (see figure 2A). In two of these models, the excitatory and inhibitory neurons were connected to generate pyramidal-interneuron network gamma (PING) oscillations^62^, with one model generating a “slow” gamma (∼32Hz) oscillation and the other generating a “fast” gamma (∼65Hz) oscillation (see figure 2B). In the remaining model, the inhibitory-to-excitatory connections were severed, preventing the generation of endogenous gamma oscillations. In this arrhythmic network, the remaining neuron parameters matched those of the “slow” gamma model, meaning this arrhythmic network had the propensity to resonate at “slow” gamma frequencies if it received rhythmic input. To explore how these models fit the behavioural data, each of these models was stimulated with 32.5Hz, 43.3Hz and 65Hz sine waves and the change in endogenous oscillatory strength was fitted to memory performance.

**Figure 2.**
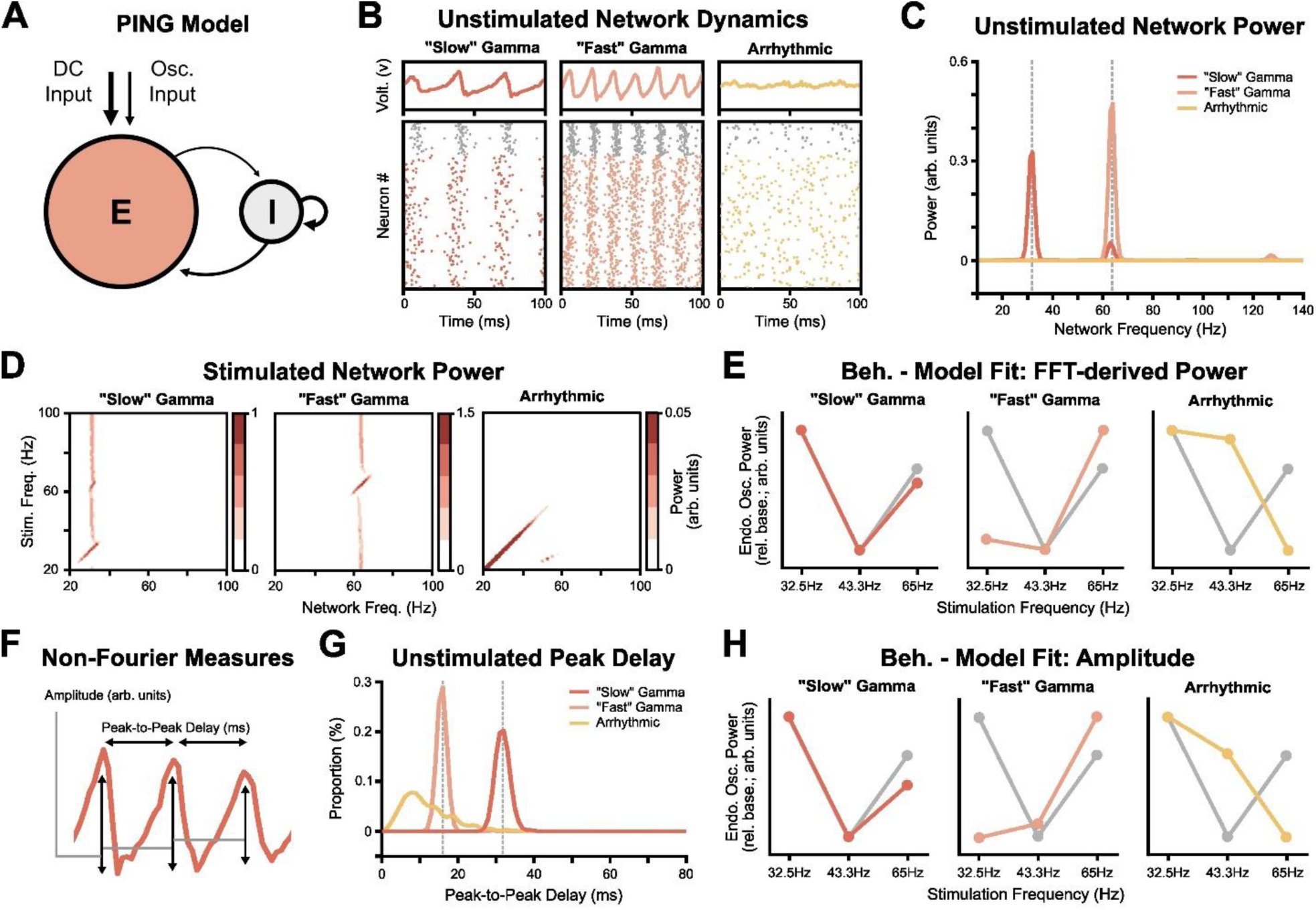
Computational modelling suggests that 65Hz and 32.5Hz sensory stimulation both entrain an endogenous 32.5Hz rhythm. **(A)** Depiction of the pyramidal-interneuron network gamma (PING) model. 320 excitatory neurons and 80 inhibitory neurons were connected to form (following direct current [DC] input) an endogenously-oscillating network. Additional oscillatory input was presented to the excitatory neurons to mimic sensory stimulation used in the behavioural experiments. **(B)** A snapshot of network dynamics for each model. The top plots reflect the mean ‘v’ parameter across excitatory neurons, which approximates the voltage from a neuron. The raster plots beneath depict firing patterns for excitatory and inhibitory neurons (in colour and in grey, respectively). **(C)** Power spectrum of excitatory neuron voltage for each model prior to stimulation. The “slow” gamma model produces a peak at ∼32.5Hz while the “fast” gamma model produces a peak ∼65Hz. No peak is observed in the arrhythmic model. **(D)** Frequency-frequency plots depicting the impact of linearly-spaced stimulation frequencies (y-axis) on the power of excitatory neuron voltage. Stimulation only modulates activity in the oscillating models when the stimulation frequency nears the fundamental or harmonic frequency of the network. **(E)** Line plots comparing group-level t-values of the behavioural data (in grey) and power of the peak frequency of computational models (in colour) following stimulation, for each model. To account for differences in scale between the behavioural data and the computational models, the measures have been scaled such that the maximum value of the behavioural data and each model equals one, and the minimum value behavioural/model data equals zero. **(F)** Depiction of the non-Fourier-based measures used to replicate the Fourier-based findings displayed in panels C-E. **(E)** Histograms of peak-to-peak delay of the excitatory neuron LFP for each model prior to stimulation. The “slow” gamma model produces a peak at ∼33ms (i.e., ∼32.5Hz) while the “fast” gamma model produces a peak at ∼15ms (i.e., ∼65Hz). The arrhythmic model peaks earlier (∼9ms) but substantially less consistently. **(H)** Line plots comparing group-level t-values of the behavioural data (in grey) and amplitude of LFP peaks (in colour) following stimulation, for each model. To account for differences in scale between the behavioural data and the computational models, the measures have been scaled such that the maximum value of the behavioural data and each model equals one, and the minimum value behavioural/model data equals zero.

In the first instance, we explored how stimulation at a linearly-spaced range of frequencies impacted the ongoing rhythms of each model. We observed that, when the stimulation frequency closely matched the fundamental frequency of the endogenous rhythm, oscillatory power increases and the endogenous rhythm shifts speed to match the stimulation frequency (see figure 2D). Critically, a similar, albeit smaller, effect can be seen when stimulating at the first harmonic frequency of the model, suggesting harmonic stimulation frequencies can modulate endogenous rhythms. No similar effect was seen for subharmonic stimulation frequencies. The arrhythmic model did not display any distinct responses to 32.5Hz/65Hz stimulation, and instead displayed a progressive decrease in oscillatory power as the stimulation frequency increased. Altogether, these findings suggest that endogenous rhythms can be entrained by harmonic stimulation frequencies (i.e., 65Hz stimulation can entrain a 32.5Hz rhythm), but not by subharmonic stimulation frequencies (i.e., 32.5Hz stimulation cannot entrain a 65Hz rhythm).

We then set out to see whether these harmonic entrainment effects could explain the behavioural results. To this end, we calculated the change in endogenous oscillatory strength following stimulation (relative to non-stimulated oscillatory strength) for each of the three stimulation frequencies and each of the three PING models. We then conducted a multiple linear regression where each PING model was used as a separate predictor regressor that aimed to explain the behavioural results reported in Experiment 1. In doing so, we found that the “slow” gamma model explained a significant proportion of variance (β = 0.416 [95% CI: 0.047, 0.785], p = 0.028; see figure 2E]. No effect was observed for the “fast” gamma model (β = 0.001 [95% CI: - 0.173, 0.175], p > 0.5), nor the arrhythmic model (β = 0.019 [95% CI: -0.177, 0.214], p > 0.5). Altogether, these results suggest that the behavioural patterns reported above are best explained by the entrainment of an ongoing endogenous “slow” gamma oscillation.

Notably, prior to stimulation, the Fourier-based decomposition of the “slow” gamma model produced a small, additional peak at ∼65Hz (see Figure 2C). While this is most probably an artifact that is generated when using a Fourier-based method on a non-sinusoidal oscillation, we nonetheless wanted to reproduce the effects using alternative, non-Fourier methods (see Figure 2F). We estimated oscillatory frequency using peak-to-peak delay and found that the distance between peaks for the “fast” gamma model were ∼15ms (i.e., ∼66Hz), distance between peaks for the “slow” gamma model were ∼32ms (i.e., ∼31Hz), and distance between peaks for the arrhythmic model were ∼9ms (i.e., ∼111Hz; though much more variable; see Figure 2G). Importantly, there was no secondary peak at ∼15ms for the “slow” gamma model, suggesting that the 65Hz peak in the FFT-derived power spectrum was an artifact of the Fourier-based decomposition as opposed to a legitimate 65Hz oscillation in the “slow” gamma model. When fitting the amplitude of the local field potential peaks to the behaviour data, we found that the “slow” gamma model continued to explain a significant proportion of variance (β = 0.118 [95% CI: 0.021, 0.215], p = 0.018; see figure 2H]. No effect was observed for the “fast” gamma model (β = 0.055 [95% CI: -0.035, 0.144], p = 0.232). The arrhythmic model also explained a significant proportion of variance (β = -0.387 [95% CI: - 0.569, -0.205], p < 0.001), but relied upon a negative beta weight to do so, suggesting the arrhythmic pattern only matched the behavioural data when the results were inverted. These results support those uncovered when using Fourier-based methods: namely, that the behavioural data is best explained by the “slow” gamma model.

We then set out to explain between-participant variability in the behavioural effects. Given that endogenous oscillatory frequencies vary between individuals, we postulated that between-participant variability in the behavioural effects could be (to some extent) explained by variability in the distance between a participant’s endogenous rhythm and the stimulation frequency. The ability of a stimulation frequency to modulate an endogenous rhythm decreases as the distance between the stimulation frequency and endogenous frequency increases (known as “Arnold Tongue”) and may also occur when stimulating at harmonic frequencies ^57–60^. To this end, we varied the endogenous frequency of the “slow” gamma model such that it ranged from 26.5Hz to 38.5Hz and then applied 32.5Hz, 43.3Hz and 65Hz stimulation. We observed the traditional Arnold Tongue, with endogenous oscillatory power increasing as the distance between the endogenous and 32.5Hz stimulation frequency decreased (see figure 3A). Critically, we also observed a similar effect when applying 65Hz stimulation, demonstrating a harmonic Arnold Tongue. Lastly, we conducted pairwise correlations of endogenous oscillatory power following stimulation between the three frequencies. Aligning with the behavioural results, we found a correlation between endogenous oscillatory power following 32.5Hz and 65Hz stimulation (r = 0.50, p = 0.014; see figure 3B), and no effect when correlating either of these frequencies with the 43.3Hz condition (32.5-43.3Hz correlation: Pearson’s r = 0.08, p > 0.5; 65-43.3Hz correlation: Pearson’s r = 0.11, p > 0.5), suggesting that the distance between the stimulation and endogenous frequency can explain between-participant variability in the behavioural responses to sensory stimulation.

**Figure 3.**
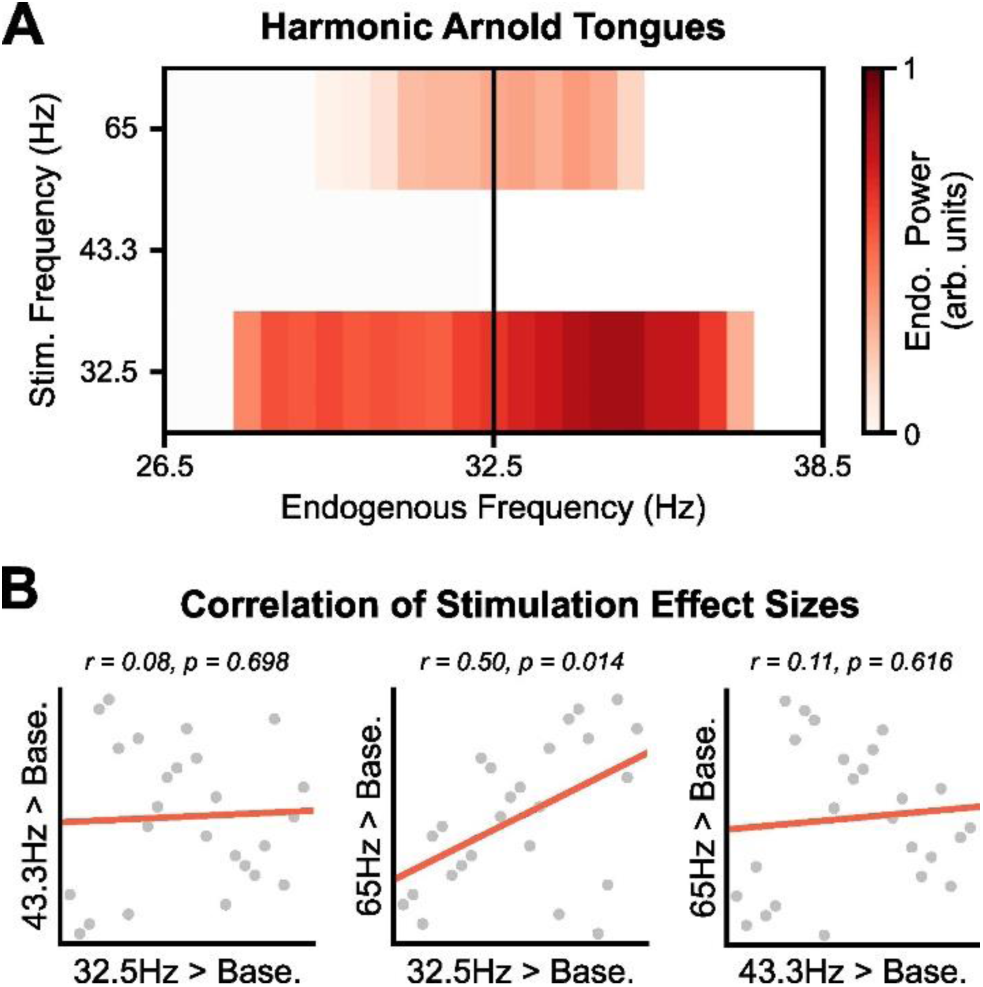
Harmonic Arnold Tongues explain between-participant variability in behavioural performance. **(A)** Frequency-frequency plot depicting how a change in endogenous oscillatory frequency impacts responsiveness to stimulation at 32.5Hz, 43.3Hz and 65Hz. The stimulation-induced change in endogenous power decreases as the endogenous frequency strays further from 32.5Hz (i.e., an Arnold Tongue). Importantly, this effect occurs for stimulation at the fundamental (i.e., 32.5Hz) and harmonic (i.e., 65Hz) frequencies. **(B)** Pairwise correlations of stimulation-induced change in endogenous power for the three stimulation frequencies. The strength of the response to stimulation at the fundamental and harmonic frequencies correlated, mirroring the correlation of behavioural effects displayed in figure 1C.

Lastly, we examined whether harmonic entrainment effects generalise beyond the first harmonic. That is, could higher (e.g., second, third) harmonic frequencies have an impact on an endogenous oscillation? To address this, we stimulated the “slow” gamma and “fast” gamma models with sine waves that approximated the first to third harmonics and the first to third subharmonics and then compared how oscillatory power following stimulation compared to a baseline measure of oscillatory power prior to stimulation. Interestingly, oscillatory power only increased above baseline when stimulating using the fundamental and first harmonic frequency (see figure 4A). This pattern of results was observed in both the “slow” gamma or “fast” gamma models. When inspecting the local field potential (see figure 4B), it appeared that stimulation using subharmonic frequencies led to mass synchronisation but impaired the periodicity of the endogenous oscillation. In contrast, stimulating at harmonic frequencies greater than the first harmonic led to a rather desynchronised network. Only stimulation at the fundamental and first harmonic frequencies produced regular, synchronised activity. Taken together, these results suggest that harmonic entrainment is only effective at the fundamental and first harmonic frequencies which, in turn, suggests that the observed behavioural effects cannot be attributed to 32.5Hz and 65Hz stimulation both entraining a slower (e.g., 16.25Hz) or faster (e.g., 130Hz) rhythm. These results further support the conclusion that 32.5Hz and 65Hz stimulation are entraining an endogenous “slow” gamma rhythm.

**Figure 4.**
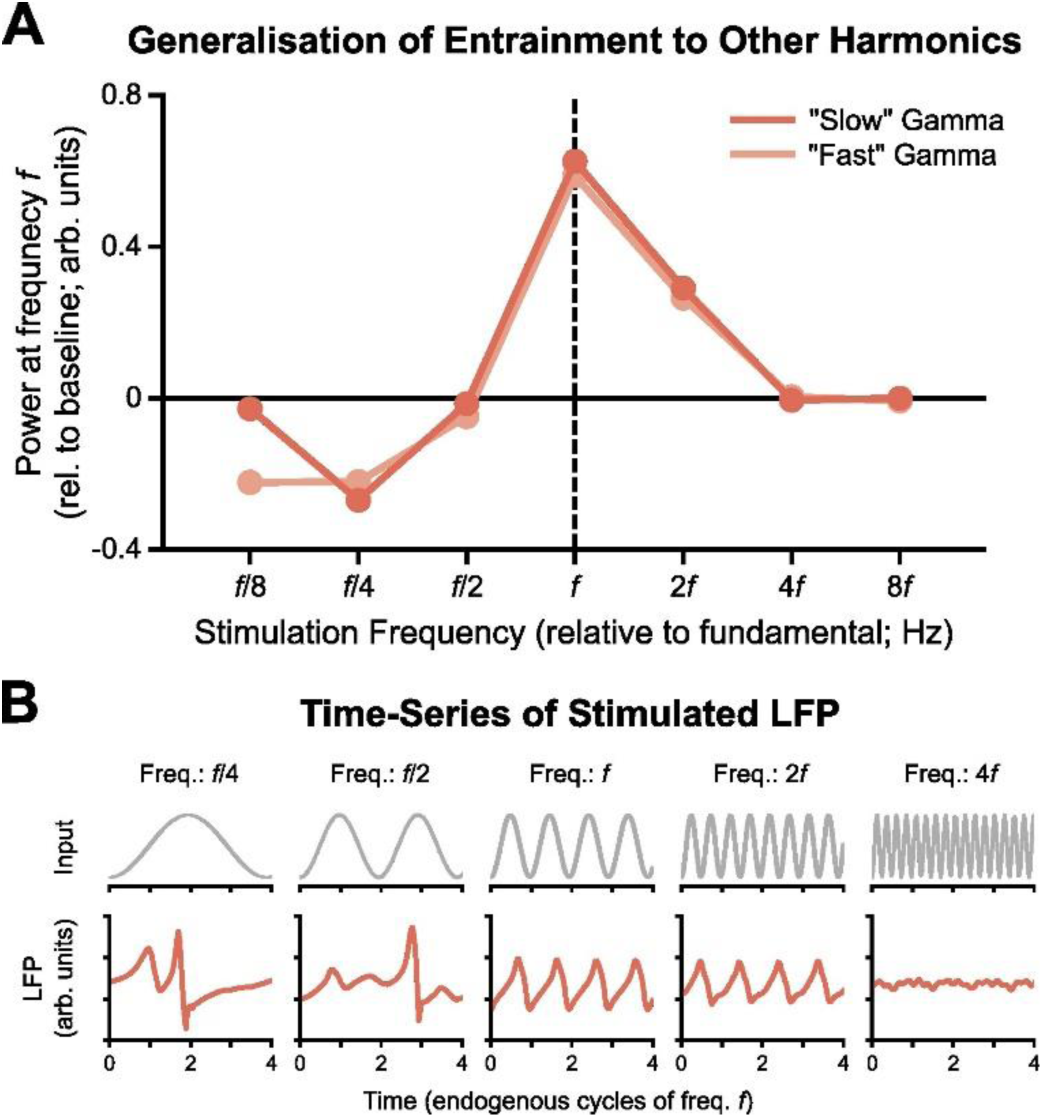
Harmonic entrainment is only effective with the first harmonic. **(A)** Line plot depicts how endogenous oscillatory power changes relative to baseline (y-axis), given stimulation frequency relative to the fundamental (f) [x-axis]. Entrainment appears to only be effective when stimulating with the fundamental or first harmonic frequency. **(B)** Time-series of local field potential following different stimulation inputs. Stimulating at the fundamental (*f*) and first harmonic (2*f*) frequencies maintain a stable endogenous gamma oscillation (see figure 2B for reference). Subharmonic stimulation (*f*/2, *f*/4) leads to mass, irregular synchronisation while stimulation at the second harmonic (4*f*) appears to desynchronise the network.

## Discussion

Here, we demonstrate that gamma-band sensory stimulation acts as an imperceptible intervention that enhances recall. When participants were presented with a retrieval cue, rhythmic fluctuations in screen luminance at 32.5Hz and 65Hz led to a statistically significant increase in the number of paired associates recalled relative to a baseline condition containing no luminance fluctuation. Critically, participants were not able to detect the 65Hz luminance fluctuation. These effects were replicated in an independent sample. Taken together, it seems that 65Hz visual sensory stimulation can be used as an intervention to aid in the retrieval of episodic memories. Additional behavioural analyses and computational modelling suggest that 65Hz stimulation operates by harmonically entraining an endogenous “slow” gamma (∼32.5Hz) oscillation. In the remainder of this paper, we discuss why the observed memory enhancement may have arisen, and how this knowledge could be utilised.

Rather than act upon an endogenous “fast” gamma rhythm, this imperceptible flicker appears to enhance recall by harmonically entraining an endogenous, “slow” gamma rhythm. This aligns with computational models and electrophysiological studies which have demonstrated that entraining at the first harmonic of an endogenous oscillation can enhance the power of that oscillation ^57–60^. Expanding on these results, our model suggests that harmonic entrainment is specific to stimulation at the first harmonic frequency. This observation aids inference as it rules out the possibility that the stimulation protocol could harmonically entrain any number of subharmonic endogenous rhythms (e.g., 16Hz, 8Hz, 4Hz). Critically, we observe harmonic stimulation impacting behaviour. This challenges the orthodoxy that behavioural changes resulting from stimulating at a given frequency are attributable to changes in endogenous activity at that same frequency. Indeed, our results suggest that in an experiment where behaviour is modulated by (for example) 8Hz stimulation, but not 6Hz or 10Hz, one cannot conclude that the modulation in behaviour is due to modulation of endogenous 8Hz rhythms. Rather, stimulation at either the harmonic (16Hz) or the subharmonic (4Hz) is required to rule out the possibility that 8Hz is harmonically entraining a 4Hz rhythm. While an additional stimulation condition may seem onerous, there is a benefit to this: our “arrhythmic” model demonstrates that harmonic stimulation effects only occur when there is an existing endogenous oscillation to entrain, meaning that the addition of a harmonic stimulation condition can allow entrainment to be inferred in the absence of electrophysiological recordings (for existing electrophysiologically-based methods that quantify entrainment, see ^63^). While further work is required to see how harmonic entrainment generalises to other forms of oscillation and other types of behavioural task, we postulate that harmonic stimulation can expedite research into neural entrainment by sidestepping the need for electrophysiological recordings to verify said entrainment, allowing for studies to be run more efficiently in a larger number of labs than has previously been thought possible.

Given that both 32.5Hz and 65Hz sensory stimulation enhances memory, and that these frequencies can act on a common rhythm through harmonic entrainment, we propose that “slow” gamma oscillations play a causal role in episodic memory retrieval. “Slow” gamma oscillations are thought to facilitate the recall of episodic memories by allowing reactivated traces to be relayed from the hippocampus out to the neocortex for vivid reinstatement ^29, 64^. These ideas implicitly suggest a causal role for “slow” gamma oscillations in recall, but evidence supporting this is limited. Our results may fill this gap. Our results cannot be explained by 32.5Hz and 65Hz stimulation modulating an arrhythmic network, nor can they be explained by these two frequencies modulating another endogenous frequency (e.g., 65Hz, 16.25Hz). Instead, it seems the results are best explained by 32.5Hz and 65Hz stimulation both entraining a “slow” gamma (∼32.5Hz) endogenous oscillation, which consequently enhances the recall of episodic memories.

Taken together, these results suggest that recall can be enhanced when the retrieval cues are presented together with an imperceptible flicker. To date, most neuroscientific-based memory interventions involve costly equipment that can only be used by specialists (e.g., transcranial magnetic stimulation [TMS]; transcranial alternating current stimulation [tACS]), and/or are uncomfortable for participants to experience (e.g., perceptible sensory stimulation ^e.g. 53^). The imperceptible 65Hz flicker used here, however, has neither of these limitations, allowing for novel, unintrusive means to augment human memory. While the flicker is not a panacea for memory (providing a boost in performance of ∼5-8% in the current task), the boost it provides is likely to be welcomed given the comparative ease and comfort with which it can be deployed relative to brain stimulation hardware or low-frequency sensory stimulation. For example, it will be of great interest to see how this protocol can be deployed in clinical settings to tackle growing issues related to age-related cognitive decline, and in educational settings to enhance developmental learning that depends on prior knowledge (e.g., language learning).

### Open Questions and Future Directions

A growing number of studies have reported the effectiveness of 40Hz sensory stimulation in enhancing episodic memory retrieval and reducing symptoms in mnemonic-related diseases (e.g., Alzheimer’s disease) ^65–67^. However, rather than deliver stimulation during recall as done here, these studies deliver stimulation for extended periods (e.g., 1 hour) during wakeful rest. Due to the difference in when these protocols are delivered, together with the absence of an effect at 43.3Hz in our experiment (the closest neighbour to 40Hz used elsewhere), we view our approach to stimulation as acting on a distinct mechanism from that of “offline” 40Hz stimulation. Such a hypothesis would suggest that the memory-enhancing effects that follow offline 40Hz stimulation are distinct from those following online 32.5/65Hz stimulation. Consequently, pairing offline 40Hz stimulation with online 32.5Hz/65Hz stimulation may enhance recall to a level greater than what is achieved using either protocol in isolation.

It remains an open question as to which brain regions are impacted by 32.5/65Hz stimulation. One candidate region is the hippocampus, for which a correlative link between memory retrieval and “slow” gamma oscillations have previously been reported^7, 29, 47, 48, 56^. These hippocampal “slow” gamma oscillations are thought to enhance connectivity between CA3 and CA1 and, consequently, relay reactivated traces from CA3 back towards the cortex. Therefore, if 32.5/65Hz sensory stimulation is enhancing hippocampal “slow” gamma oscillations, it would suggest that our stimulation is specifically aiding the reactivation process. However, it is important to note that there is an ongoing debate about whether sensory stimulation can reach the hippocampus. Intracranial recordings in rodents and MEG recordings in humans suggest that visual flickers are unlikely to reach the hippocampus ^51–53^. However, this may be because the subjects were not cognitively engaged: when subjects are engaged in tasks with working/episodic memory components, gamma-band sensory stimulation does appear reach the hippocampus^54, 55^ (though these observations were made with source-localised EEG, which struggle to accurately measure the hippocampus). In short, one could argue that 32.5/65Hz sensory stimulation aids recall by boosting hippocampal “slow” gamma activity, but further investigation that blends spatial specificity (e.g., intracranial rodent recordings) with cognitive engagement (e.g., an episodic memory paradigm) is required to verify whether these flickers can indeed reach the hippocampus.

There is also the question as to why stimulation was ineffective during encoding. Numerous previous studies have linked enhanced “fast” gamma oscillatory activity to successful memory formation, but our attempts to modulate this rhythm during encoding did not influence subsequent memory. Given the effectiveness of stimulation during retrieval, it cannot be said that 65Hz stimulation simply does not impact ongoing neural activity. However, it may be that 65Hz stimulation cannot impact patterns of gamma activity relevant to memory formation. For example, decreases in alpha power have been reported to precede and predict increases in gamma power during episodic memory formation ^29^, meaning the effectiveness of 65Hz stimulation on encoding could be mediated by these alpha oscillations. Based on these ideas, it seems premature to conclude that “fast” gamma oscillations do not play a causal role in episodic memory formation. Instead, additional future studies are required to rule out the possibility that confounding variables are masking the link between 65Hz sensory stimulation and episodic memory formation.

### Conclusion

In sum, we demonstrate that imperceptible, high-frequency sensory stimulation can enhance mnemonic performance. While questions remain about the underlying circuits which are influenced by this protocol, it may nonetheless provide a cost-effective, unintrusive and easy-to-implement intervention to augment human memory all the while hinting at a causal role of “slow” gamma activity in episodic memory retrieval.

## Methods

### Participants

Forty-four participants took part in Experiment 1 (38 female; mean age: 23.2, age range: 18-31). After outlier removal (see below for details), 34 participants remained (29 female; mean age: 23.4, age range: 18-31). All participants were right-handed, native German speakers. Based upon an effect size of d = 0.82 (taken from a pilot study) and an alpha of 0.05, this sample had statistical power that exceed β = 0.99. All participants provided informed consent prior to taking part in the experiment. Ethical approval was granted by the ethics committee at Ludwig-Maximilians-Universität, Munich, Germany.

Thirty-seven new participants took part in Experiment 2 (26 female; mean age: 21.9, age range: 18-31; 4 left-handed). After outlier removal (see below for details), 29 participants remained (21 female; mean age: 21.8, age range: 18-31; 4 left-handed). Based upon an effect size of d = 0.57 (taken from the results of Experiment 1) and an alpha of 0.05, this sample had statistical power of approximately β = 0.85. All participants provided informed consent prior to taking part in the experiment. Ethical approval was granted by the ethics committee at Ludwig-Maximilians-Universität, Munich, Germany.

### Paired-associates task

Participants completed a video-word paired-associates task previously shown to correlate with memory-related gamma-band power changes during both encoding and retrieval 29. We presented the task on a Scaleo C792 monitor with a refresh rate of 130Hz (visual display: 20cm x 28cm [H x W]). Participants sat approximately 75cm from the screen. In Experiment 1, participants completed 3 blocks of the experiment, with each block consisting of 64 video-word pairs. In Experiment 2, participants completed 3 blocks of 48 video-word pairs. Prior to completing the main task, participants completed a brief demo consisting of four trials that acquainted them with the task.

During encoding, participants were presented with a fixation cross for approximately 1,500ms (with a uniform jitter of 200ms), followed by a video clip for three seconds (video display size: 10cm x 12cm [H x W]), and then a German noun for another three seconds (word display size: 1cm x ∼4cm [H x W]). There were four videos in total, each paired to 48 nouns (or 36 nouns for Experiment 2). After seeing the video-word pairing, participants were presented with a question. In Experiment 1, this question always asked whether the participant felt there was a semantic association between the video and the word. This question was used to keep participants focused on the stimuli, rather than provide any meaningful metric for later analysis. In Experiment 2, this question would either ask about the semantic association (as in Experiment 1) or ask whether the screen was flickering during the presentation of the stimuli. The latter question was subsequently used to explore the perceptibility of the visual flicker. The questions alternated in a pseudo-random order, such that each question was presented an equal number of times. After participants responded or if three seconds elapsed, the next trial began.

During encoding, we flickered the screen throughout the presentation of the fixation cross, video and word. In Experiment 1, the flicker was presented at either 65Hz, 43.3Hz or 32.5Hz, or not at all. The flicker was created by presenting a black box (on-screen size: 14cm x 13cm [H x W]) on top of the stimuli, occluding them from view, every second frame (for 65Hz flicker), third frame (for 43.3Hz) or fourth frame (32.5Hz) [no black box was presented for trials where no flicker was presented]. In Experiment 2, the flicker parameters remained the same but only the 65Hz or “no flicker” conditions were used.

After completing the encoding phase, participants completed a brief distractor task. Here, participants were presented with a series of simple mathematical subtractions in the centre of the screen, and were asked to type in the correct answer. If participants answered correctly, the answer turned green and a new sum appeared. If participants answered incorrectly, the answer turned red and they were asked to complete the sum again. Once two minutes had elapsed and the participant had solved at least ten problems, the retrieval task began.

During retrieval, participants were presented with a fixation cross for approximately 1,500ms (with a uniform jitter of 200ms), followed by a German noun for another three seconds. These nouns always came from the immediately-preceding encoding phase (i.e., there were no new nouns or lures from other blocks). The word then disappeared and participants were presented with stills from the four videos. Here, they were asked to select which video they thought was associated with the word. After participants responded or if three seconds elapsed, a confidence question appeared asking whether the participant felt they could remember experiencing the association, simply knew the association, or had forgotten the association. After participants responded or if three seconds elapsed, the next trial began.

During retrieval, the screen flickered throughout the presentation of the fixation cross and word and ceased at the onset of the video selection screen. The flicker parameters matched those used during encoding.

### Flicker detection task

At the end of the paired-associates task, participants who completed Experiment 1 went on to complete a short detection task in which video-word pairs were presented in the same manner as the encoding phase of the paired-associates tasks. During stimulus presentation, the screen flickered at 65Hz, 43.3Hz, or 32.5Hz, or did not flicker. At the end of the fixation-video-word sequence, participants were asked whether the screen had flickered. After participants responded (or if three seconds elapsed) the next trial began. Participants completed 120 of these flicker detection trials in a single block. Note that only 38 participants completed the flicker detection task.

### Model-based analysis of paired-associates task

While the main results of this paper focus on different hypotheses than those originally pre-registered (see https://osf.io/qkm2g/; see supplementary table 1), we nonetheless endeavoured to keep our analytical pipeline as close as possible to that which was pre-registered. Times in which we deviated from the pre-registered analysis plan have been described in the same supplementary table that presents the original pipeline.

For each participant, we first excluded any trials in which participant did not respond to the semantics/flicker question at encoding, target selection at retrieval, or confidence at retrieval as a means to exclude trials in which participants were perhaps not attending to the screen. We then derived a measure of memory performance in which “successful recall” was described as any trial in which participants chose the correct video and reported that they did not forget the association. Trials in which participants chose the incorrect video, or reported that they had forgotten the association, were recorded as “forgotten”. We then used L1-regularised logistic regression to predict this measure of memory performance across trials based upon a series of predictors:

1. “Flicker at encoding”. This reflect a series of binary regressors (one for each flicker frequency) that marked trials with the target flicker frequency at encoding (e.g., 65Hz) as 1 and all other frequencies are marked as 0. In Experiment 1, this produced three regressors (i.e., one regressor for 65Hz, one for 44.3Hz, and one for 32.5Hz). In Experiment 2, only a single regressor for 65Hz was used.
2. “Flicker at retrieval”. This reflect a series of binary regressors (one for each flicker frequency) that marked trials with the target flicker frequency at retrieval (e.g., 65Hz) as 1 and all other frequencies are marked as 0.
3. “Reaction time”. This reflected the time it took participants to identify which video was associated with the presented word. This served to control for a speed-accuracy trade-off, addressing concerns that a particular flicker frequency boosts/slows reaction times, which in turn impairs/boosts memory performance^68^.
4. “Experimental time”. This reflected two regressors (one for encoding, one for retrieval) that linearly increases in value for each trial. This regressor will capture any variation in performance that can be explained by cognitive exhaustion experienced during the course of the task.
5. “Intercept”. Modelled as a vector of ones, this served to de-mean the outcome variable and allowed us to interpreted a positive regressor coefficent for the flicker regressors as a memory-enhancing effect, and a negative coefficient for these regressors as a memory-impairing effect.

This model produced a coefficent for each of these regressors for every participant, which was then converted into a t-value by dividing the beta value by the standard error of the fit. One-sample t-tests were used to test whether the standardised regressor coefficents for a given flicker condition deviated from zero beyond what would be expected by chance. Outliers were removed prior to group inferential statistics (see “outlier removal” below). Given the exploratory nature of Experiment 1, two-tailed tests were used and multiple comparisons were corrected for using the Bonferroni method. As Experiment 2 served to replicate the findings of Experiment 1, one-tailed tests were used to explicitly test whether the 65Hz flicker had a memory-boosting effect when presented during encoding.

### Outlier removal

Outlier removal followed our pre-registered pipeline. We removed participants that fell outside the inclusion criteria in a stepwise manner, so that the outliers did not influence subsequent inclusion criteria. The stepwise procedure worked as follows:

1. Participants who had a hit-rate (i.e., the proportion of correctly-selected videos, regardless of confidence) of less than 30% were excluded. This served to limit the likelihood that someone who was simply guessing would be included in the final analysis. At this stage, no participants were removed from Experiment 1; three participants were excluded from Experiment 2.
2. Participants with memory performance (i.e., the proportion of correctly-selected videos that were not marked as “forgotten) greater than 80% or less than 20% were excluded. This served to prevent ceiling/floor effects obscuring the effects of interest. Here, an additional nine participants were removed from Experiment 1; an additional five participants were excluded from Experiment 2.
3. In the case of the participant-level logistic regression analyses, participants with a flicker regressor coefficient that exceeded three absolute deviations from the group median were marked as outliers. Here, one additional participant was removed from Experiment 1; no additional participants were excluded from Experiment 2.

### Percentage-based analysis of paired-associates task

In addition to this modelling approach, we also estimated the percentage change in successful recall for flickers relative to baseline (i.e., when no flicker was presented). The same inferential tests (i.e., one-sample t-tests) were used to probe whether the percentage change in memory performance for each flicker condition was significantly greater than zero.

### Flicker detection analysis

In experiment 1, participants completed a separate task where they had to report whether they could perceive a flicker during the presentation of video-word pairs (for details, see the section “Paired-associates task”). For each flickering frequency (32.5Hz, 43.3Hz, and 65Hz), trials where the screen was flickering were pooled with trials where no flicker was presented, resulting in 60 trials (30 with a flickering screen and 30 without a flickering screen). We computed “flicker detection accuracy” by calculating the percentage of times participants correctly reported seeing the flicker / correctly reported not seeing the flicker. To determine whether a participant could reliably detect the flicker, the labels of each trial were shuffled and the flicker detection accuracy was recomputed. This process was repeated 500 times to build a surrogate distribution describing the likelihood of detecting the flicker by chance. Participants with a flicker detection accuracy greater than 95% of the surrogate samples were said to be able to reliably perceive the flicker. The same analytical procedure was applied to the data collected in Experiment 2.

### PING model architecture

We modelled a cortical area of interconnected 320 regular spiking excitatory pyramidal neurons and 80 fast-spiking inhibitory neurons. The quantity of neurons and the ratio of excitatory-to-inhibitory neurons approximates those used in previous studies ^61, 69, 70^. The architecture of the model was built upon that proposed by Izhikevich^61^, which simulates the membrane potentials of both excitatory and inhibitory neurons. Parameter values are reported in Table 1.

**Table 1.**
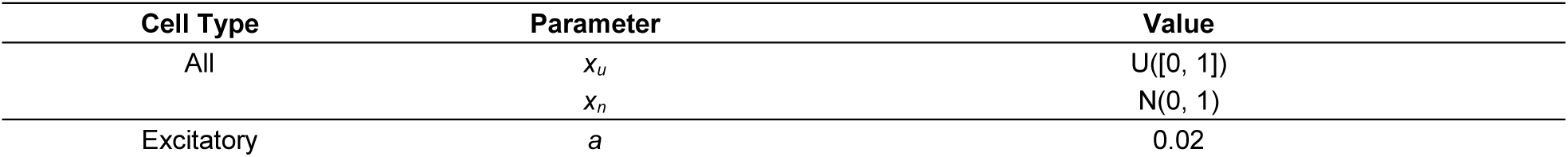
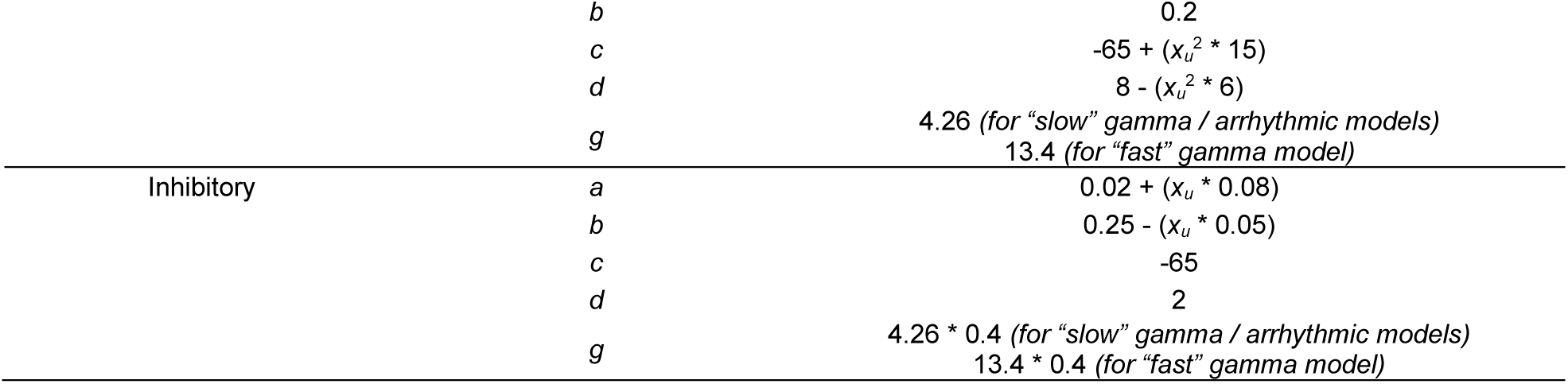
Model parameters for excitatory and inhibitory cells. All parameters used in the Izhikevich equations (Equations 1-3) match those reported in the original paper. Note that for the computing of the Arnold Tongues visualised in Figure 3, the values of *g* were systematically varied between 1.6 and 5.4, in steps of 0.02.

The equations below describe how the membrane potential of one neuron is calculated:

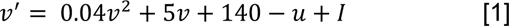

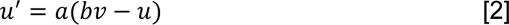

Here, *v* approximates the membrane potential of the simulated neuron, *I* reflects the input current, and *u* is a recovery variable. When *v* exceeds 30mv, and action potential is modelled:

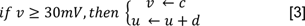

The input *I* varied based on cell type and model type. It is modelled as the sum of direct current input (*I_DC_*), input from connected cells (*I_endo_*) and a sinusoidal input (*I_exo_*).

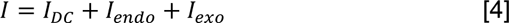

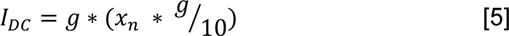

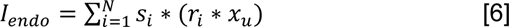

Here, *s* denotes a vector of 400 binary values (i.e., one for each neuron) that describes which neurons fired at *t_(n-1)_*; *r* denotes a vector of 400 connectivity values that describe the strength of the pairwise connections between the simulated neuron and all other neurons in the network. For inhibitory-to-excitatory connections, *r* = -0.6; for inhibitory-to-inhibitory connections, *r* = -1; for excitatory-to-inhibitory connections, *r* = 0.5; For excitatory-to-excitatory connections, *r* = 0. All connectivity values were multiplied by a random value, uniformly sampled between 0 and 1. The simulated neuron had no connection to itself. For the arrhythmic model, the inhibitory-to-inhibitory and inhibitory-to-excitatory connections were set to 0 as a means to mimic AMPA receptor suppression that results in a reduction in the amplitude of gamma oscillations ^71^.

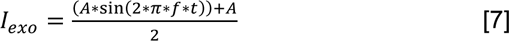

Here, *A* was set to 10% of *g* (for depictions of how this ratio impacts the model, see supplementary figure 2); *f* denotes the stimulation frequency; *t* denotes the current sample (in milliseconds). Note that equation 7 denotes a standard sinusoid equation with an additional scalar that ensures the minimum value of *I_exo_* = 0 and the maximum value of *I_exo_* = A.

### Model analysis

The remainder of the analysis of the model was conducted on the population activity of the excitatory cells (that is, the mean of the *v* parameter across all excitatory neurons), which approximates the signals recorded by macroscopic measures such as MEG.

Spectral power of the population time series was computed using a sliding-window approach to FFT^72^. The FFT was computed for windows of 1 second (sample rate = 1000Hz), with 95% overlap between neighbouring windows. For each of the three stimulation frequencies (32.5Hz, 43.3Hz, and 65Hz), power at the spectral peak following stimulation at frequency f was then contrasted against power at the spectral peak when stimulation was not applied (i.e., *I_exo_* = 0). This returned the change in peak spectral power given stimulation at frequency f (from here on referred to as “stimulation-modulated oscillatory power”).

Non-Fourier equivalents to this measure were computed to address concerns about harmonic artifacts that can arise when using Fourier-based methods to decompose non-sinusoidal methods. Peak-to-peak distance was computed by first z-scoring the time-series of the v parameter (averaged over all excitatory neurons, which approximates local field potential) and then identifying any peaks that exceed one standard deviation of the mean of the signal. Histograms were built to visualise variability in peak-to-peak delay and infer stationary rhythms (see figure 2F). The amplitude of the oscillations was quantified as the mean amplitude of all identified peaks.

To quantify how the stimulation-modulated oscillatory power and amplitude fitted to the behavioural data, a multiple regression was conducted with an outcome variable y reflecting the behavioural data (i.e., the first-level t-values of the logistic models), and three predictor variables reflecting the three computational models (*x_slow-gamma_, x_fast-gamma_, x_arrhythmic_*). As the behavioural and computational measures were in different unit spaces, they were both z-transformed prior to regression model fitting. This meant that all regressors (predictors and outcomes) had a mean of zero, circumventing the need to include a constant in the model. The “OLS” function in the statsmodel package was used to estimate model fit and significance.

## SUPPLEMENTARY MATERIALS

Supplementary figures 1-2

Supplementary table 1

Summary of results for the pre-registered hypotheses

**Supplementary Table 1.** A description of the pre-registered experimental design as if 9^th^ April 2021 (original pre-registration available at https://osf.io/qkm2g/). Any deviations from the approach are listed in the right hand-column.

| Section | Pre-registered Protocol | Deviations from Protocol |
| --- | --- | --- |
| Description | Episodic memory is highly dependent on the hippocampus, and the neural oscillations present within. Recent work in humans (Griffiths et al., 2019, PNAS) has demonstrated that episodic memory formation is linked to hippocampal fast gamma activity (~65Hz), while episodic memory retrieval is linked to hippocampal slow gamma (~45Hz) activity. However, these links are strictly correlative. Here, we aim to address this central limitation by using visual entrainment (at fast and slow gamma frequencies) to influence episodic memory formation/retrieval and hence demonstrate a causal link between fast/slow gamma and episodic memory formation/retrieval |  |
| Hypotheses | 1) Flickering stimuli at 65Hz ("fast" gamma) during encoding should see greater cued recall performance at retrieval when compared to the aggregate of all other conditions.<br>2) Flickering stimuli at 43.3Hz ("slow" gamma) during retrieval should see greater cued recall performance at retrieval when compared to the aggregate of all other conditions. |  |
| Study Type | Experiment - A researcher randomly assigns treatments to study subjects, this includes field or lab experiments. This is also known as an intervention experiment and includes randomized controlled trials |  |
| Blinding | No blinding is involved in this study. |  |
| Is there any additional blinding in this study? | No blinding is involved in this study. Participants may perceive differences in flickering rate but will be unaware of the hypothesis. The experimenter will observe the participant from another room and can't see the presentation monitor, hence they cannot influence participant behaviour in line with the hypotheses. |  |
| Study-design | Within-subjects design. |  |
| Randomisation | Participants will be shown pseudo-randomised flickering stimuli. The flickering will be randomised such that every combination (e.g. 65Hz at encoding and 43.3Hz at retrieval; 43.3Hz at encoding and 43.3Hz at retrieval etc...) occurs equally often. |  |
| Existing Data | Registration prior to analysis of the data |  |
| Reason for existing data | Data for 10 participants has already been collected. The data of 4 of these participants was accessed to check that the files were being written correctly. No analyses of any type were conducted. |  |
| Data collection procedures | Participants will receive course credit (1 credit per hour) in return for their time. Participants must be between 18 and 35 years of age. Participants must be native German speakers (as some stimuli are German words). | Some participants received financial payment in place of course credit |
| Sample size | 32 human participants |  |
| Sample size rationale | An a priori power analysis was conducted using a pilot sample. Given alpha = 0.05, power = 0.95, and Cohen's d = 0.82 (derived from pilot sample), 32 participants would be the required sample to observe the hypothesised effect. |  |
| Stopping rule | Data collection will cease once 32 datasets have been collected. Outlier analysis will then be conducted (see below), and outlying participants will be removed from the sample. Additional participants will then be recruited to reach the sample size target, and outlier analysis will once again be conducted. This process will continue iteratively until the sample target is reached. |  |
| Manipulated variables | We will manipulate the frequency at which stimuli at encoding and at retrieval flicker (32.5Hz, 43.3Hz, 65Hz, no flickering) on a trial-by-trial basis. The flicker frequency of a pair at encoding is independent of the flicker frequency of that same pair at retrieval. |  |
| Measured variables | We will measure the number of pairs that are correctly recalled, and the response time to select the associated stimulus. |  |
| Indices | Recall performance will be binary (they recalled the pair, or they didn't). Reaction times will be measured in seconds from when selections could be made. |  |
| Statistical models | For each subject, a logistic regression will be run where memory performance (recall vs. forgotten) will be predicted by four variables ("flicker at encoding", "flicker at retrieval", "reaction time", and "linear").<br>1) "Flicker at encoding" will be a binary regressor where the 65Hz ("fast" gamma flicker) at encoding is marked as 1 and the other frequencies are marked as 0.<br>2) "Flicker at retrieval" will be a binary regressor where the 43.3Hz ("slow" gamma flicker) at retrieval is marked as 1 and the other frequencies are marked as 0.<br>3) "Reaction time" will be a continuous regressor that marks the time taken to select a stimulus. This controls for speed-accuracy trade-off, addressing concerns that a particular flicker frequency boosts/slows reaction times, which in turn impairs/boosts memory performance.<br>4) "Fatigue" will be a continuous regressor bound between 0 and 1, that linearly increases in value for each trial. This regressor will capture any variation in performance that can be explained by cognitive exhaustion during the course of the task.<br>This approach returns four coefficients (two of interest; i.e. flicker frequency) for each subject. The two coefficients of interest will be contrasted in separate, one-tailed, one-sample t-tests. These tests would examine whether the coefficients of the sample are greater than zero (i.e. flickering at the hypothesised frequency boosts memory performance). | L1 regularisation was applied to the logistic regression to enhance model stability. This decision was made during the analysis of the data for Experiment 1, and prior to the collection of data for Experiment 2.<br>The "fatigue" regressor was split into two regressors. One describing trial number at encoding, and the other describing trial number at retrieval (the original pre-registered approach did not account for the fact that stimulus order at retrieval was shuffled relative to the encoding order). As with the L1 regularisation, this decision was made during the analysis of Experiment 1, and prior to the collection of data for Experiment 2. |
| Transformations | Recall performance on each trial will be coded as "hit" or "miss". A hit refers to a stimulus pair that was correctly recalled, and not marked as a 'guess' when probed for the 'remember/know/guess' judgement. A miss refers to a stimulus pair that was not correctly recalled, or was marked as a 'guess' when probed for the 'remember/know/guess' judgement. |  |
| Inference criteria | We will use the standard $p < 0.05$ criteria for determining whether the t-tests reveal any differences between conditions. | |
| Data exclusion | 1) Trials where no response to stimulus, familiarity, or semantics was recorded will be removed from analysis. This reduces the likelihood of including a trial where a participant was not attending to the screen/task.<br>2) Participants who select the correct stimulus during the task less than 30% of the time will be excluded (ignoring |  |
|  | the remember/know/guess judgement), as they are likely to be exhibiting chance performance (chance = 25%).<br>3) Participants who recall more than 80% hits or less than 20% hits will be excluded to reduce the risk of floor/ceiling effects masking any results.<br>4) Participants with coefficients that deviates from the group (as measured by absolute deviation around the median; Leys et al., 2013, J. Experimental Social Psychology) will be excluded [median +/- 3 times the MAD method]. |  |
| Missing data | Participants who failed to complete the experiment will be excluded from all analysis. |  |
| Exploratory analysis | An additional logistic regression will be conducted on recalled trials to test whether flicker frequencies influence reaction time. The linear regression and statistical tests will be conducted in the same manner as above. Moreover, another linear regression will test whether 43.3Hz flickering at encoding and 65Hz flickering at retrieval is detrimental to memory performance. The regression will be conducted in a similar manner to what is described above, but the “flicker at encoding” binary regressor refers to 43.3Hz and the “flicker at retrieval” binary regressor refers to 65Hz. These analyses are complementary to the key hypotheses, and may prove interesting supplements to the main result. |  |

### Summary of results for the pre-registered hypotheses

Previous work in humans has revealed two distinct hippocampal gamma oscillations that correlate with episodic memory: a “fast” ∼65Hz gamma that relates to encoding, and a “slow” ∼45Hz gamma that relates to retrieval (e.g., Griffiths et al., 2019). Here, we set out to test whether these correlative effects reflect a causal relationship in which “fast” and “slow” gamma aid the formation and retrieval of memories respectively. To this end, we used visual sensory stimulation while participants encoded and retrieved video-word pair associations, and explored how the frequency of the visual stimulation affected recall performance.

The analyses were conducted on the data obtained from Experiment 1, and conducted in a similar manner to what was reported in the main text. For each participant, a L1-regularised logistic regression was used to predict memory performance based on the presence of 65Hz visual stimulation at encoding, 43.3Hz visual stimulation at retrieval, reaction time, and fatigue. The standardised regressor coefficients were pooled together across participants, and one-sample t-tests then compared whether these coefficients deviated from zero.

Matching what was reported in the main text, no effect of “fast” gamma was observed on memory formation [t(34) = -0.11, p > 0.5]. However, a significant decrease in memory performance occurred when “slow” gamma sensory stimulation was applied during memory retrieval [t(34) = - 3.14, p = 0.003]. To explore what drove memory detriment effect, a second model was conducted, where each flicker frequency was modelled separately (note: this is the model reported in the main text). Based on the nature of this model, we are able to observe how memory performance during visual stimulation varies purely from memory performance in the absence of visual stimulation. Here, the detrimental effect of “slow” gamma disappeared, and was replaced by significant 32.5/65Hz effects reported in the main text. This suggests that the original detrimental effect of “slow” gamma during retrieval was most probably due to large beneficial effects from the 65/32.5Hz visual stimulation, rather than a meaningful detrimental effect for 43Hz.

The results presented here find no evidence to suggest that “fast” gamma sensory stimulation can influence memory formation, nor can ∼45Hz gamma sensory stimulation influence memory retrieval. Rather, it seems that a slower gamma rhythm (∼32.5Hz) is key to recall. While there is a frequency mismatch between the previously-reported ∼45Hz gamma effect (Griffiths et al., 2019) and the 32.5Hz effect observed in the main text, the 32.5Hz effect is nonetheless aligned with the “slow” gamma reported in rodents (e.g., Colgin, 2009; Zheng et al., 2016). While this mismatch occurred remains unclear, though may possibly be due to the fact that the existing human recordings of “slow” gamma were conducted in patients with epilepsy, who may have atypical endogenous rhythms.

**Supplementary Figure 1.**
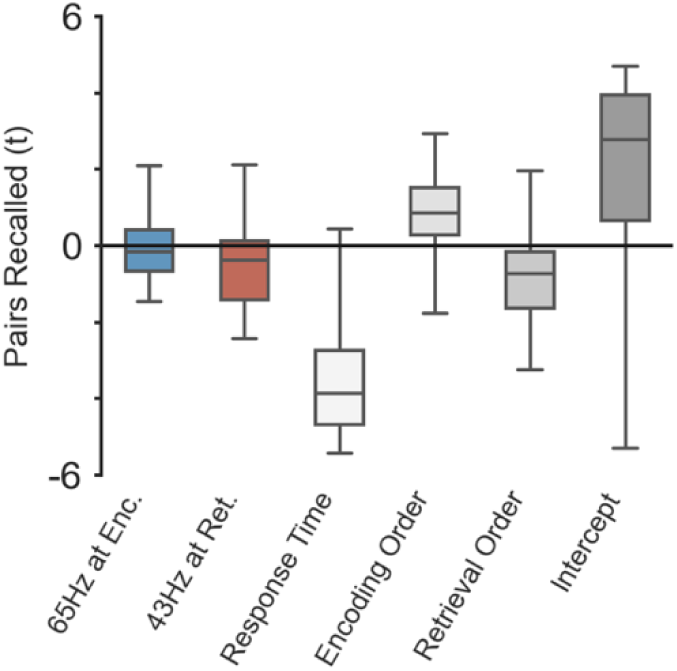
Boxplot depicting impact of defined predictor variables on the number of pairs that were successfully recalled. Each boxplot reflects a separate regressor included in the pre-registered model. The central line of each boxplot reflects the median, the bounds of each boxplot reflect the 25^th^ and 75^th^ quartiles, and the tails reflect the smallest and largest participant-specific values. While we hypothesised a memory-enhancing effect for the two flicker conditions (coloured boxplots), no significant effect was found. Instead, a significant decrease was uncovered for the 43Hz flicker during retrieval (though this transpired to be an illusionary effect driven by large positive effects for 65Hz and 32.5Hz retrieval flickers).

